# Simulating homeostatic, allostatic and goal-directed forms of interoceptive control using Active Inference

**DOI:** 10.1101/2021.02.16.431365

**Authors:** Alexander Tschantz, Laura Barca, Domenico Maisto, Christopher L. Buckley, Anil K. Seth, Giovanni Pezzulo

**Author notes:** Corresponding author: Giovanni Pezzulo ISTC-CNR, Via San Martino della Battaglia 44 00185 Rome, Italy.

## Abstract

The adaptive regulation of bodily and interoceptive parameters, such as body temperature, thirst and hunger is a central problem for any biological organism. Here, we present a series of simulations using the framework of Active Inference to formally characterize interoceptive control and some of its dysfunctions. We start from the premise that the goal of interoceptive control is to minimize a discrepancy between expected and actual interoceptive sensations (i.e., a prediction error or free energy). Importantly, living organisms can achieve this goal by using various forms of interoceptive control: homeostatic, allostatic and goal-directed. We provide a computationally-guided analysis of these different forms of interoceptive control, by showing that they correspond to distinct generative models within Active Inference. Furthermore, we illustrate how these generative models may support empirical research, by predicting physiological and brain signals that may accompany both adaptive and maladaptive interoceptive control.

**Highlights:**

- We use Active Inference to provide formal models of interoceptive control
- We model homeostatic, allostatic and goal-directed forms of interoceptive control
- Our simulations illustrate both adaptive interoceptive control and its dysfunctions
- We discuss how the models can aid empirical research on interoception

## Introduction

Living organisms constantly face adaptive regulation problems, such as keeping bodily and interoceptive variables such as temperature and glucose levels within acceptable ranges, in the face of external hazards. The solution of these biological problems is so fundamental for our existence that it plausibly shaped the design of our brains, much before we were able of sophisticated forms of cognition [1]; and this is well testified by the centrality of interoceptive circuits in neuronal architectures [2–4].

From a formal perspective, the adaptive regulation of bodily and interoceptive parameters is challenging, and benefits from predictive (allostatic) forms of control. The theory of allostasis emphasizes that to regulate bodily and interoceptive parameters efficiently, it is important to anticipate bodily needs (e.g., future increases of body temperature or oxygen demands) and prepare to satisfy them before they arise (e.g., transpiration or hyperventilation) rather than just react to sensed bodily or interoceptive dysregulations [5].

Mechanisms of predictive or allostatic control in living organisms can vary greatly in their complexity. In our previous example, a predicted increase of body temperature or oxygen demand directly triggers autonomic processes for transpiration or hyperventilation. This is a rather simple form of predictive control, which only engages autonomic and interoceptive systems. More sophisticated ways to solve the same regulation problems go beyond autonomic reflexes of this kind. For example, advanced organisms (like humans) can counteract the increase of their body temperature by buying and drinking a glass of wine or by finding shade on a hot sunny day. From a formal perspective, engaging these complex goal-directed loops requires sophisticated cognitive architectures that are multimodal, hierarchical and/or temporally deep [6]. Multimodality is important when it is necessary to coordinate exteroceptive and interoceptive streams; for example, to predict that some gustatory stimuli (e.g., those sensed when drinking wine) can eventually solve an interoceptive problem (e.g., thirst). Temporal hierarchy is important when the phenomena of interest span different timescales. For example, obtaining and drinking a glass of wine involves a sequence of actions (a ‘compound action’) that lasts seconds or minutes, whereas interoceptive reflexes are much faster, taking place on the order of hundreds of milliseconds. Finally, temporal depth refers to the hierarchical depth of the planning process. It becomes important, for example, when a living organism has to predict how its actions now will change its interoceptive streams in the future; for example, the fact that carrying a bottle of water in hot sunny day is useful, even if one is not currently thirsty. These features of multimodality, temporal sequencing and temporal depth pose additional challenges and opportunities for more complex forms of predictive allostatic control.

These challenges can be usefully articulated within a formal approach to adaptive regulation and allostasis provided by the perspective of Active Inference: a leading theory in computational neuroscience [7,8]. Active Inference describes the brains of living organisms as “prediction machines” that solve biological regulation problems by learning internal (generative) models of their bodily and interoceptive processes [9] and of how they can produce desired outcomes by acting in the world [10,11]. It appeals to the normative imperative of prediction error (more formally, free energy) minimization, to explain a broad range of cognitive and emotional processes, such as perception and attention [12–14], goal-directed control [8,15,16], interoceptive processing [11,17–19] and various other aspects of cognitive processing [20–22]. Furthermore, Active Inference has been recently used to shed light on various psychopathological conditions, such as psychosis, depression, eating disorders and panic disorders, in terms of aberrant inference [23–30].

Here, we use Active Inference to develop four simulations of interoceptive control. Our simulations cover different forms of control, which range from the simple correction of sensed interoceptive errors to the more complex proactive regulation of the system that renders it able to anticipate and prospectively regulate future interoceptive errors. Furthermore, our simulations allow the exploration of both adaptive and maladaptive cases of interoceptive control, of the kind associated to psychopathologies [31].

One strength of Active Inference is that modelling these different forms of interoceptive control does not require changing the inferential framework. Rather, it only requires expanding the simulated organism’s generative model, by including more modalities (e.g., only interoceptive or also exteroceptive modalities) and hierarchical levels (e.g., only fast-changing or also slowly-changing variables). Another strength of Active Inference is that dysregulations of interoceptive control (that are increasingly recognized as central to understanding various forms of psychopathology) emerge naturally from suboptimal settings of the model parameters [32]. Our simulations illustrate these strengths by showcasing how varying critical model parameters (e.g., the precision or inverse variance of interoceptive streams) can render simulated organisms more or less sensitive to interoceptive sensations and consequently, more or less able to achieve allostasis. Finally, following a theory of valence formulated within Active Inference [33], we will simulate the affective responses of simulated organisms to changes of their interoceptive variables, hence providing a formal link between the realms of body regulation and affect.

Our simulations offer both theoretical and methodological contributions. First, they provide a computationally-guided theoretical analysis of various forms (uni-vs. multimodal; shallow vs. hierarchical) of interoceptive control, from the normative perspective of Active Inference. Second, they illustrate (simulated) physiological and brain signals that one should expect to measure in biological organisms that engage in adaptive or maladaptive (e.g., psychopathological) forms of interoceptive control. The simulated signals are important as they can be used for model-based data analysis (or “computational phenotyping”) in future studies of interoceptive processing in both healthy individuals and those suffering from interoceptively-mediated psychopathologies [34].

## Methods

Active Inference is a theory from computational neuroscience that describes perception, action and learning within a unified framework [7,8]. It proposes that the brain forms a generative model of its sensory data, which comes to encode knowledge about the environment’s statistical contingencies. On this account, perception is the process of “inverting” this model to infer the environment’s state from noisy and ambiguous sensory data. Similarly, learning can be described as the process of inverting a model which includes beliefs about parameters [35]. Finally, Active Inference can account for behaviour in terms of inverting a model which encodes beliefs about actions (proprioceptive variables).

Under Active Inference, the inversion of generative models is achieved through variational inference, an optimisation procedure which approximates Bayesian inference. To implement this optimisation, Active Inference suggests that the brain’s internal states (e.g., neural activity) parameterise an approximate posterior distribution, which describes an agent’s beliefs about the (hidden) state of the world. These internal states then change in order to minimise an information-theoretic quantity, the variational free energy *F*, which quantifies the Kullback-Leibler divergence (KL-divergence) between the approximate posterior and the generative model:

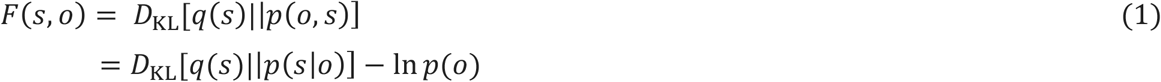

where *q* (*s*) is the approximate posterior over hidden states *s*, *p* (*o*, *s*) is the generative model over observations *o* and hidden states *s*, and *D*_KL_ is the KL-divergence. The second equality demonstrates that variational inference minimises the KL-divergence between the approximate and true posterior distributions, thereby providing an approximation to Bayesian inference [8,36].

Active Inference extends this approximate inference scheme to account for action. Mathematically, it posits an approximate posterior over states *and* actions *q* (*s*, *a*), and a generative model over observations, states and actions *p*(*o*, *s*, *a*). Actions are then inferred through the minimisation of variational free energy, with the generative model specifying the prior probability of actions. To account for purposeful, goal-driven behaviour, Active Inference proposes that an agent’s generative model assigns a higher probability to preferred (e.g., rewarding) outcomes. This move ensures that variational free energy is minimised when sampling desired outcomes (e.g., receiving a reward). As behaviour can change which outcomes are sampled, minimising variational free energy with respect to action entails sampling preferable parts of the environment (e.g., searching for a reward). On this view, perception and learning correspond to updating one’s beliefs to match the environment, whereas action corresponds to sampling the environment to match one’s beliefs.

### Predictive coding

While Active Inference provides a general scheme for describing perception, action and learning, the specific updates depend on the types of distribution employed. In computational neuroscience, a popular update scheme is provided by predictive coding, which treats both the approximate posterior and generative model as Gaussian distributions [12,37]. Under these assumptions, variational free energy can be rewritten in terms of precision weighted prediction errors:

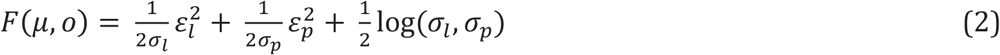

where *ε_l_* = *o* − *g*(*μ*) and 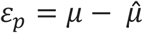 are *prediction errors, μ* is the mean of the approximate posterior *q* (*s*), *g*(*μ*) is some (possibly non-linear) function describing the relationship between states and observations, 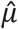 is the mean of the prior distribution over states *p* (*s*), and *σ* are the variances of the corresponding distributions [12,36]. (Note that the term ‘precision’ here refers to inverse variance.)

A prediction error describes the difference between a prediction and the actual input, whereas the variance corresponds to the uncertainty in the prediction (or the input). Precision weighted prediction errors are formulated in terms of differences in mean values weighted by their relative precisions. Predictive coding is usually considered in the context of hierarchically organised generative models, where each level predicts the activity of the level below it (besides the lowest level, which predicts sensory data). In this context, beliefs and model parameters at each level are updated to minimise Equation 2, and the resulting scheme can be straightforwardly implemented in biologically plausible networks composed of prediction units (which convey predictions) and error units (which convey the difference between predictions and inputs) [38]. The network dynamics ensure that prediction errors are minimised over time, such that predictions and parameters come to capture the data in a more accurate manner.

In recent years, this predictive coding scheme has been generalised to action [8,39]. Here, actions are deployed to minimise proprioceptive prediction errors (e.g., a classical reflex arc), where predictions about proprioceptive signals are provided by hierarchically superordinate layers. Crucially, these actions can still be cast as performing gradient descent on free energy, but now with respect to sensory data rather than beliefs. Therefore, while perception minimises prediction error through an iterative updating of descending predictions, action minimises prediction error by directly changing the sensory input. To incorporate some notion of value or desire, an agent’s generative model can be equipped with a set of prior beliefs which are consistent with that agent’s continued survival. For instance, a model may encode the belief that the average body temperature will be around 37 degrees. Significant deviations from these prior beliefs will result in prediction errors which are then minimised through action. This process model implements both interoceptive inference, where autonomic actions minimize interoceptive prediction errors [2,9,11,18] and goal-directed behavior, where externally-directed actions minimize proprioceptive (and exteroceptive) prediction errors [39,40].

### Partially observable Markov Decision Processes

Most implementations of predictive coding assume a discrete updating scheme operating over continuous variables. These implementations are naturally suited to many biologically relevant quantities, such levels of light or degree of muscle extension. However, organisms also make decisions that involve selecting among a set of discrete actions or plans. To accommodate scenarios like this, Active Inference has been extended in the context of partially observable Markov decisions processes (POMDPs), in which the model variables have discrete values and change at discrete time intervals rather than on a continuous basis. In POMDPs, the approximate posterior and generative model are both described by categorical distributions (with Dirichlet priors) [8,41]. In the same manner as predictive coding, perception and learning are achieved through minimising variational free energy. However, the POMDP formulation supports planning, and the evaluation of entire courses of actions (or policies) rather than just the next action. Formally, this implies that actions are now selected based on the minimizing the *expected free energy G*(*π*), which quantifies the free energy that is expected to occur from executing some sequence of actions or policy *π*:

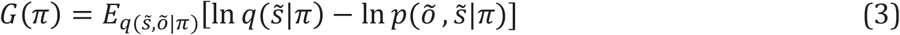

where the ~ notation denotes a sequence of variables over time. Expected free energy scores policies in terms of their expected consequences, and as such enables prospective, goal-directed action. It can be decomposed into instrumental and epistemic value terms, where the former quantifies the degree to which expected observations correspond with prior beliefs (e.g., preferences), and the latter quantifies the amount of information an agent expects to gain from some policy. See [8,42] for technical account of expected free energy.

### Hybrid models

Motivated by the fact that brains must deal with both continuous and discrete quantities, recent work has proposed a hybrid approach that utilises both types of model to describe perception, action and learning [41]. On this account, hierarchically shallow layers are implemented using predictive coding over continuous variables, as these layers directly interface with continuous sensory data and output continuous actions. In contrast, hierarchically deeper utilize POMDPs, as these layers evaluate and decide between a discrete set of action policies.

In such hybrid schemes, in order to enact a selected policy, it is necessary to map from discrete policies (e.g., picking up a bottle of water) to continuous commands (e.g., contract muscles by a certain amount). To achieve this mapping, the outcomes of the POMDP are treated as priors for the subordinate predictive coding model. These prior beliefs define the set point for the predictive model, which are then realised through action. Formally, this mapping is implemented as a Bayesian model average over the predicted outcomes from a POMDP, see [41]. While the discrete model constrains the continuous model’s activity, the continuous model provides observations or evidence for the discrete model. In the simulations presented here, these ascending messages are implemented by discretising the output from the top layer of the continuous model after it has converged.

## Results

This section illustrates four simulations of interoceptive control (and its disorders) using increasingly complex generative models within an Active Inference scheme. The first simulation illustrates a simple scenario involving homeostatic regulation through autonomic reflexes. The second incorporates anticipatory, allostatic control by coupling exteroceptive sensations to beliefs about future interoceptive states. The third simulation explores how aberrant precision weighting can lead to maladaptive regulation. Finally, the fourth simulation introduces a hybrid model combining high-level discrete policy choices with the low-level continuous predictive coding schemes as in the first three simulations. This fourth simulation enables goal-directed allostatic regulation.

### First simulation: reactive (homeostatic) interoceptive control

The first simulation illustrates a simple example of reactive (homeostatic) interoceptive inference: the case of a living organism who keeps an interoceptive variable under control in spite of external disturbances. For illustrative purposes, we will refer to this interoceptive variable as “body temperature” in this and the following simulations. However, please note that the models discussed below are generic and not intended to mimic the biological details of thermoregulation, which are significantly more complex than discussed here [43]. Rather, our goal is to illustrate the broad principles of how an Active Inference approach can be used to model interoceptive inference and (mal)adaptive regulation.

In Active Inference terms, the preferred ‘target’ value of the interoceptive variable is encoded as a prior, analogous to a “set point” in cybernetics [44–46]. Any significant deviation from the prior produces an interoceptive prediction error, which has to be minimized. This first simulation illustrates the physiological and brain signals that one should observe when a homeostatic challenge (e.g., excessively high body temperature) arises that elicits an interoceptive prediction error, which in turn triggers autonomic reflexes (e.g., vasodilatation) that respond to and suppress this error. Note that the prediction error can be conceptualized in Bayesian terms as a “surprise”, in the sense of a deviation from an expected value; and is calculated in Active Inference as a “free energy” (see the Methods section). For simplicity, we will use these terms as synonyms in this and the next simulations. Finally, this simulation illustrates the possible emotional valence associated to interoceptive regulation. For this, we formulate valence as the negative rate of free energy over time [33]. In keeping, with this formulation in the current simulation, valence is negative as free energy increases (e.g., when some external event causes a sudden increase of body temperature, beyond the desired value) and positive when free energy is being minimised (e.g., when an autonomic reflex restores the desired temperature value). This aspect of the simulation is intended to mimic a “readout” of a person’s positive and negative feelings during interoceptive regulation.

The generative model used in the first simulation is shown in Figure 1A. This model is endowed with the prior belief (denoted *μ_prior_*) that interoceptive sensory data (body temperature) will remain at 0 degrees (we use 0 degrees for mathematical simplicity – you may think of 0 degrees as corresponding to a more normal thermal set-point of ~37 degrees if helpful). In turn, these prior beliefs predict posterior beliefs about body temperature (*μ_intero_*), which themselves predict the interoceptive sensory data which conveys the current body temperature (*y_intero_*). The states and observations are treated as continuous variables and updated according to the predictive coding scheme described in the Methods section. When the body temperature deviates from prior beliefs, interoceptive predictions errors arise, due to the fact that posterior beliefs cannot satisfy both constraints simultaneously. However, these prediction errors can be minimised at the level of autonomic reflexes (*a*), which minimise the errors by updating sensory data so that it conforms to beliefs. In the current simulations, autonomic reflexes can directly update interoceptive sensory data, meaning that the gradient descent scheme for action can be specified in terms of interoceptive prediction errors:

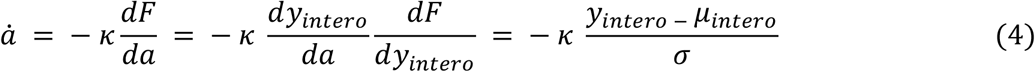

where *k* is the learning rate.

**Figure 1.**
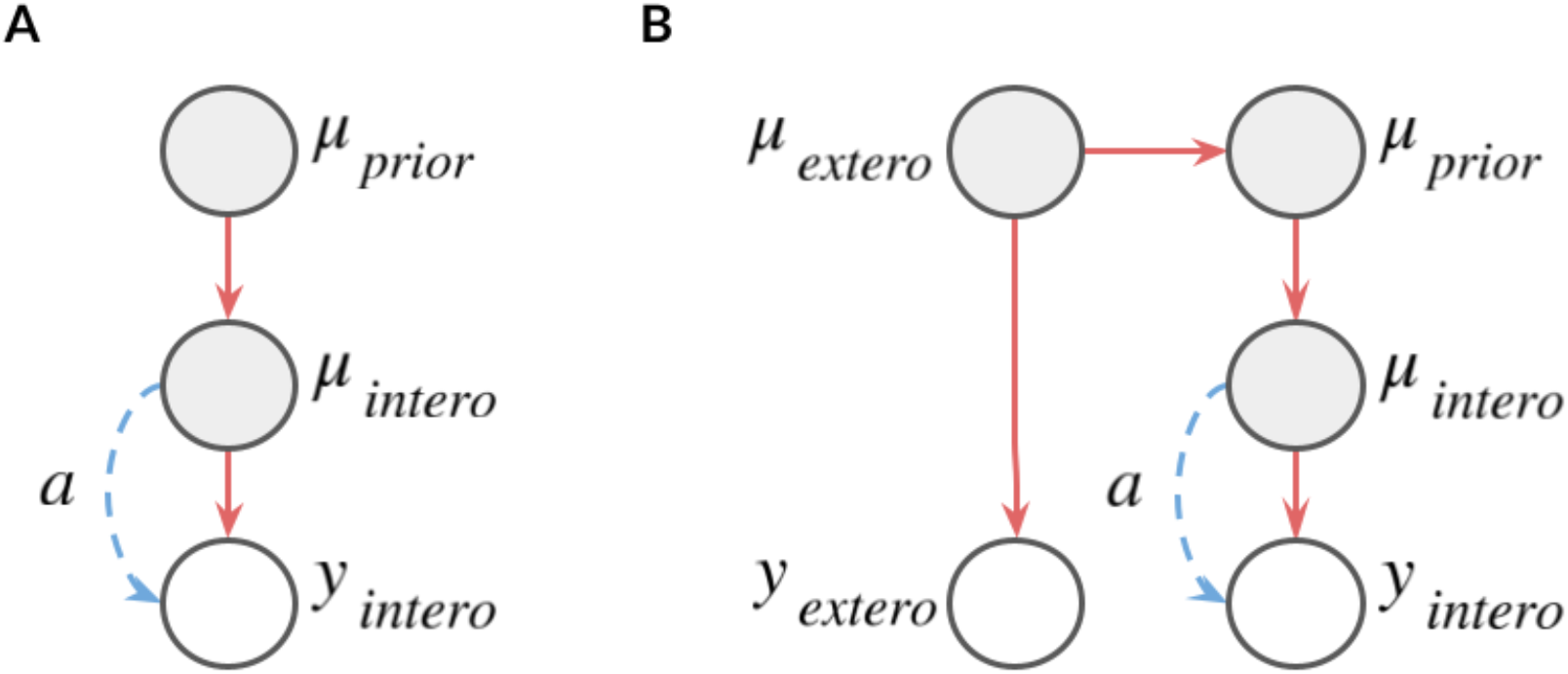
Generative models used in the first and second simulations. (A) Generative model used in the first simulation. Mu prior is the prior belief that body temperature will remain at 0 degrees. Mu intero is the posterior belief about body temperature, which is inferred based on the prior and the interoceptive data (y intero). Autonomic reflexes (a) can cancel out any prediction error, which arises from a deviation between the prior belief and the sensed body temperature. (B) Generative model used in the second simulation. The model is the same as the first simulation, but it also includes a belief about an exteroceptive signal (mu extero) and the sensed external signal (y extero). See the main text for explanation.

The results of the first stimulation are shown in Figure 2. The six panels of the figure show the values of the variables of the generative model of Figure 1A (plus auxiliary variables), over time. These are the body temperature (*intero data*, Figure 2A), the internal representation of body temperature (*Mu intero*, Figure 2B), the prior or “set point” of body temperature (*Mu prior*, Figure 2D), the autonomic action (Figure 2E), free energy (Figure 2C) and valence signal (Figure 2F).

**Figure 2.**
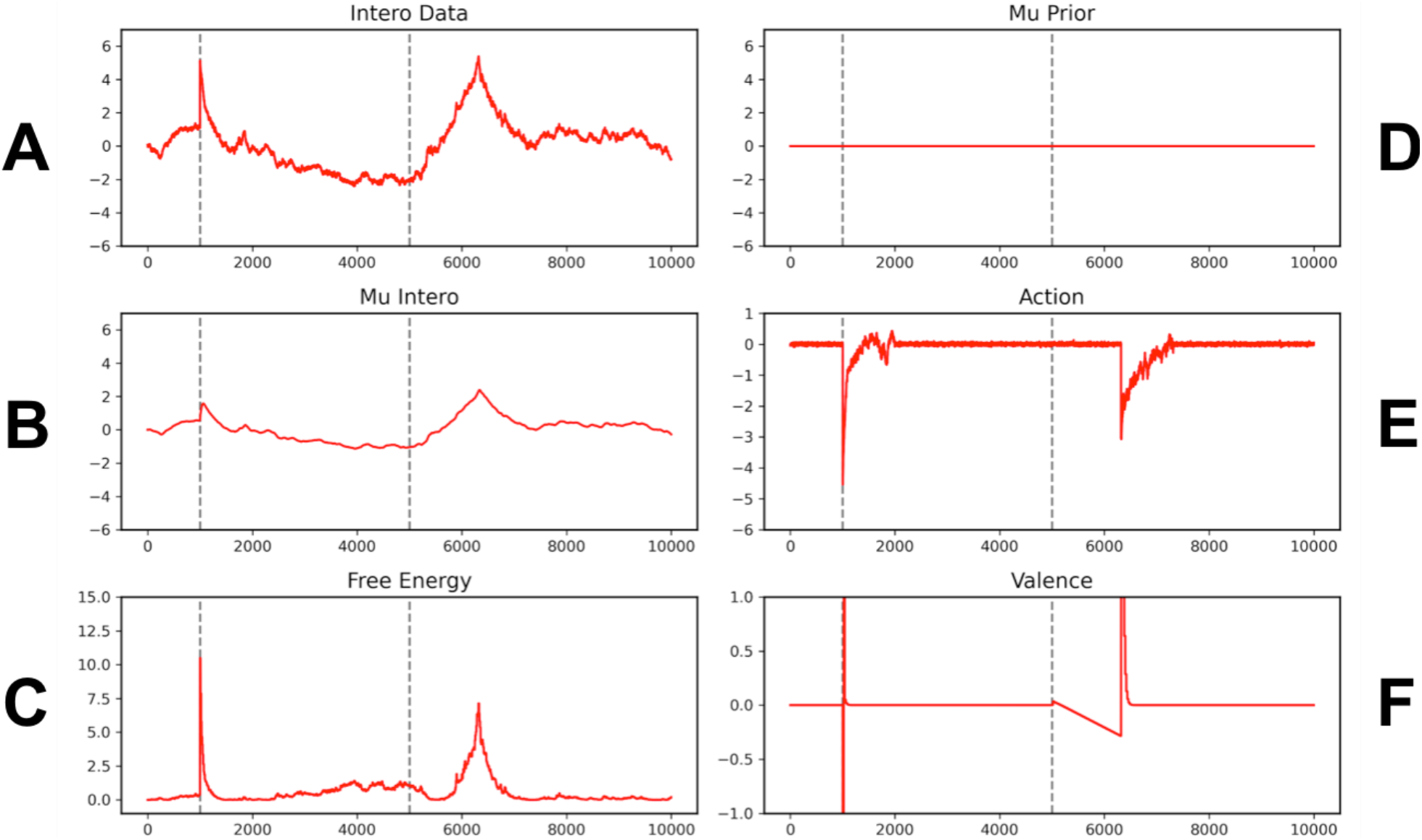
Results of the first simulation. The six panels show the values of the variables of the generative model of Figure 1A (and auxiliary variables), over time. (A) Interoceptive (body temperature) sensations. Temperature is shown in the ordinate and time is shown in the abscissa (in this and the following panels, units are arbitrary). (B) Central representation of (posterior belief about) body temperature. (C) Free energy as a measure of interoceptive error (i.e., discrepancy between prior and posterior beliefs about body temperature plus the discrepancy between posterior beliefs and data). (D) Prior over body temperature, here fixed to 0 for all time. (E) Autonomic action that is elicited to compensate for the increases of free energy shown Figure 2C. (F) Current valence, as measured by the negative rate of change of free energy. See the main text for explanation.

At the beginning of the simulation, body temperature (top-left panel) starts from a homeostatically acceptable value (here, 0 degrees). However, during the simulation an external event (e.g., a hot sunny day) determines two rapid increases of body temperature that perturb homeostasis. These two increases are indicated by the two vertical bars: the former, more rapid increase occurs at time point 1000 and the latter, slower increase starts at time point 5000. As a consequence, the internal representation of temperature (Figure 2A) also changes over time. Note that the internal representation is not the same as the interoceptive inputs, but rather a “compromise” between the prior and the inputs. This can be appreciated by noticing that the time series of the Mu intero variable has the same shape as the time series of the intero Data, but its numerical values are significantly scaled down. The deviations of the posterior belief about body temperature from the sensory data are registered as “surprise" signals (or more formally, increases of free energy, bottom-right panel), which trigger autonomic actions (Figure 2E) that cancel them out, hence restoring homeostasis.

As shown in Figure 2F, interoceptive changes that are behaviorally relevant can engender a phenomenological counterpart: they can be associated to positive or negative feelings (positive or negative valence). In keeping with [33], we calculated valence as the negative rate of free energy over time, such as it is negative when surprising deviations from homeostasis occur and positive when autonomic actions restore homeostasis. Specifically, the simulation shows that the steep increases of free energy around times 1000 and 5000 will be felt as negative events at the phenomenological level (e.g., distress); but the subsequent restorations of homeostasis will be felt as positive events (e.g., relief). While the notion of valence used in these simulations is simplified and the magnitude of the positive and negative events is arbitrary, these results illustrate possible ways the information-theoretic concept of free energy can help bridge the gap between interoceptive regulation and emotional cognition [11,25].

This first simulation illustrates the functioning of a simple generative model of Active Inference for homeostasis, which only includes two hidden variables and is unimodal (i.e., it only models interoceptive streams). The two variables are the prior and a posterior belief about body temperature, respectively. The former is fixed (here, it is zero) and represents the desired homeostatic value; whereas the latter is an estimate of interoceptive state, which combines prior information and interoceptive sensations. When the sensory data and prior beliefs disagree, the posterior beliefs converge to a midpoint between these incompatible sources of information. The discrepancy between posterior beliefs and sensory data causes a “prediction error”, which triggers an autonomic action that cancels it out. Of course, given that the prior is fixed, the only way the autonomic reflex can cancel out the error is by changing interoceptive signals, e.g., decrease body temperature (with vasodilatation) if its value exceeded zero. The possibility to act upon sensations is what distinguishes Active Inference from predictive coding, which can reduce errors only by changing beliefs. Finally, this simulation illustrated the potential readout of positive and negative feelings during interoceptive inference (in this simple simulation, these valence values are simply ‘read out’, but are not used during inference). This last aspect of the simulation can help explore the well established connection between bodily regulation and emotional valence (e.g., the positive or negative character of emotion) [11,25].

The homeostatic model used here roughly corresponds to early cybernetic control schemes, such as the TOTE model of [47]. Despite its simplicity, this model can correct (at least some) unpredicted deviations from homeostasis, by minimizing free energy. However, this generative model is limited and only supports reactive control: it can only compensate for sensed deviations from the (fixed) prior, but it cannot anticipate deviations and prepare to deal with them in advance. Furthermore, the only way it can cancel out interoceptive prediction error is by engaging autonomic reflexes. In the next simulations, we will illustrate more sophisticated generative models of Active Inference that relax these constraints and afford more effective forms of adaptive regulation.

### Second simulation: predictive (allostatic) interoceptive control

The second simulation extends the first simulation and illustrates predictive/anticipatory (allostatic) interoceptive inference. Imagine the case of an experienced runner, who participates very often in running competitions. For the athlete, participating in a competition produces a predictable increase of body demands (e.g., oxygen or glucose) and associated interoceptive changes (e.g., body temperature and ventilation), which in some cases can be very significant. In these conditions, reactive (homeostatic) strategies that attempt to maintain a fixed prior belief about the interoceptive variable of interest (e.g., body temperature) can be ineffective. For example, if deviations are too large, reactive control can fail to restore homeostasis in an acceptable time [5].

Instead, the theory of allostasis proposes that the brain exploits the predictability of (some) bodily and interoceptive changes. A way to deal with a predictable increase of bodily demands is to make prospective changes to the prior belief about the interoceptive variable of interest. From a physiological perspective, making these prospective changes may correspond to mobilizing bodily resources (e.g., oxygen or glucose levels) in advance, as often observed in athletes before a competition. While this advance mobilization may disturb homeostasis in the short run, it can help the organism deal more effectively with future necessities and compensate future external perturbations more effectively, hence reducing average prediction error (or free energy) in the long run.

In this simulation, we illustrate the allostatic regulation of body temperature in an experienced runner before and during a competition. As remarked above, here thermoregulation is used only for illustrative purposes; the model is generic and not intended to mimic the biological details of the thermoregulatory system.

The generative model used in the second simulation is shown in Figure 1B. This is an extended (multimodal) version of the generative model used in the first simulation, which also includes an exteroceptive variable that is informative about a future increase of body temperature (e.g., a visual signal indicating that the competition is about to start). Crucially, in the agent’s generative model, the exteroceptive and interoceptive variable are coupled, such that the appearance of the exteroceptive stimulus (the visual signal) functions as a cue that body temperature is about to increase, hence permitting the agent to take corrective actions in advance.

More formally, in comparison to the Figure 1A, the generative model shown in Figure 1B includes an additional exteroceptive modality *y_extero_*. The model estimates the posterior distribution over this modality through *μ_extero_*, which can be interpreted the agent’s beliefs about the causes of exteroceptive sensations. Moreover, this model encodes knowledge that exteroceptive and interoceptive data are related (e.g., that running leads to an increased body temperature and increased metabolic cost). This relation is formalised by having exteroceptive beliefs predict (e.g., act as a prior on) interoceptive beliefs. Specifically, the model encodes an inverse relationship between these two variables: as exteroceptive beliefs rise (fall), the model predicts that interoceptive beliefs will fall (rise) by an equal amount. While the fact that the relationship is inverse may sound strange, the reason is that the prior belief about body temperature in this model represents a desired value or set point, not a belief in the standard sense. What the agent wants is avoiding an excessively high body temperature. For this, when a cue that the body temperature is about to increase appears, it has to down-regulate the prior or set point, so that an autonomic action (e.g., vasodilatation) can be engaged that keeps body temperature low ^1^. Interesting, in this simulation the autonomic action is engaged before an actual change of body temperature is sensed, hence preventing body temperature from reaching excessively high values. This means that exteroceptive cues can give rise to a proactive change in autonomic processing, in the absence of any interoceptive changes.

The results of the second stimulation are shown in Figure 3. Similar to Figure 2, this figure shows interoceptive data (Figure 3A), Mu intero (Figure 3B), autonomic action (Figure 3C), free energy (Figure 3D), Mu prior (Figure 3G) and valence (Figure 3H). However, it also includes two additional panels, showing exteroceptive inputs (Figure 3E) and their internal representation (Figure 3F).

**Figure 3.**
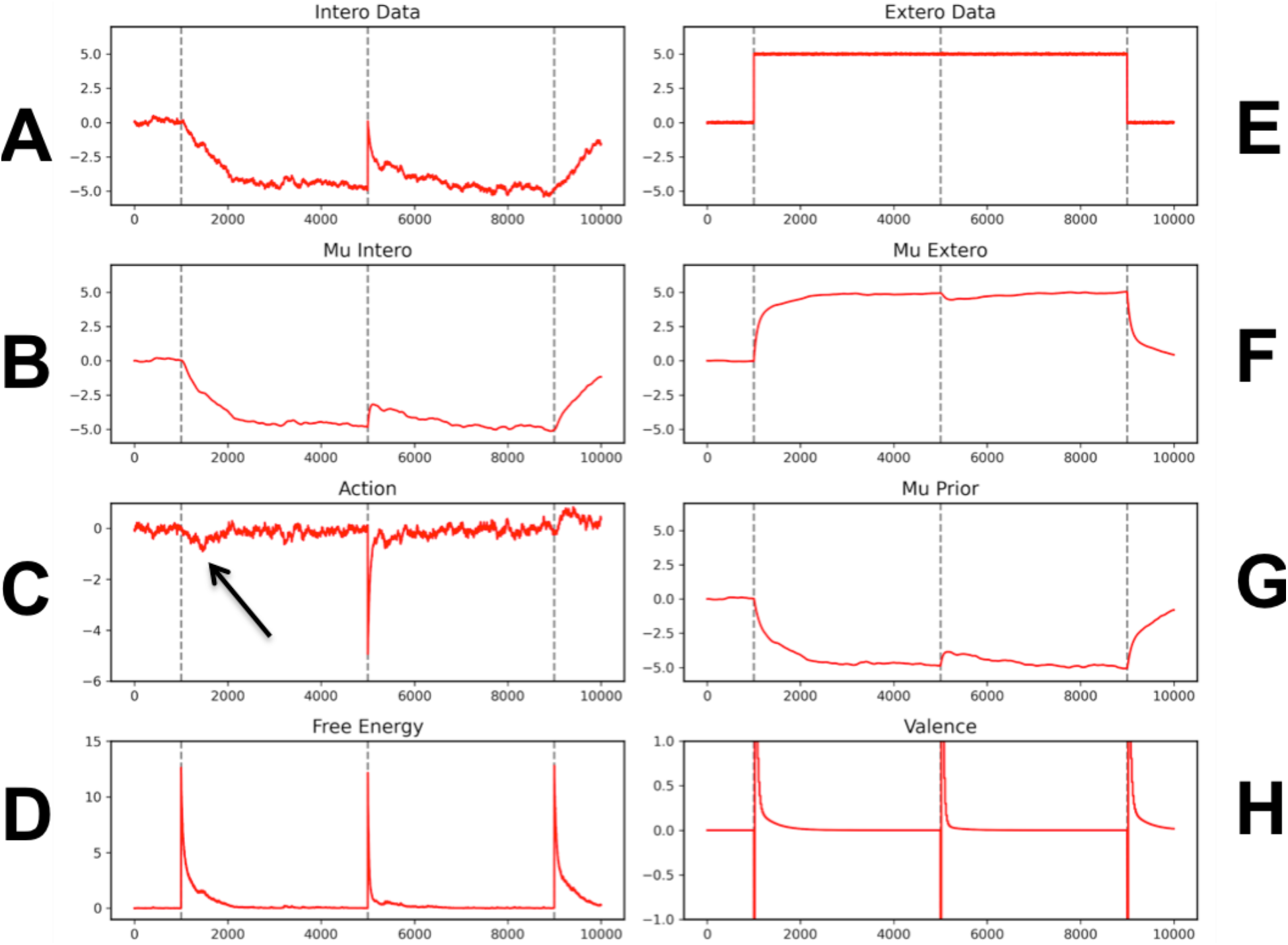
Results of the second simulation. (A) Interoceptive (body temperature) sensations (B) Central representation of (posterior belief about) body temperature. (C) Autonomic action that is elicited to compensate for the increases of free energy shown in Figure 3D. (D) Free energy as a measure of interoceptive error (i.e., discrepancy between prior and posterior beliefs about body temperature). (E) Exteroceptive sensations. (F) Internal representation (posterior belief about) exteroceptive sensations. (G) Prior over body temperature. (H) Current valence, as measured by the negative rate of change of free energy. See the main text for explanation.

In this simulation, the first vertical bar indicates the beginning of the exteroceptive signal (e.g., a signal that the competition is about to start), the second vertical bar indicates an increase of body temperature (here, we simulated a sudden change for simplicity) and the third vertical bar indicates the end of the exteroceptive signal (e.g., end of competition).

In the simulation, the exteroceptive signal (first vertical bar) causes a sudden decrease of the prior belief over temperature, which in turn causes a small but important autonomic action (see the black arrow in Figure 3C) that acts upon the sensed body temperature (Figure 2A). It is possible to interpret this autonomic action as a proactive vasodilatation that causes body temperature to (slightly) decrease, to prevent the possibility that future increases of body temperature during the run will bring this variable outside acceptable bounds. This anticipatory strategy is effective: during the run, when the actual increase of body temperature occurs (second vertical bar) body temperature does not exceed zero. With a purely reactive (homeostatic) controller, the increase of temperature would have been significantly greater.

It is worth noting that in the first simulation, all the significant changes to peripheral signals (body temperature, Figure 2A), central interoceptive representations (Mu prior, Figure 2D; and Mu intero, Figure 2B) and valence (Figure 2F) were triggered by external events that act upon the interoceptive system (i.e., at the times indicated by the two vertical bars). By contrast, in this second simulation these variables change also as an effect of internal adjustments before any actual change of interoceptive inputs. For example, the decrease of body temperature apparent after the first vertical bar is due to the internal down-regulation of the prior (that triggers an autonomic response), not to an external cause.

This proactive element of the simulation highlights important methodological implications for experimental studies of interoception. During such experimental studies, it is common to manipulate external variables (e.g., show emotional images), which are treated as “stimuli” or “inputs” to the system of interest, in order to measure peripheral and central “responses” or “outputs”. However, this simulation illustrates that the input-output approach is insufficient to model a system whose behaviour is not captured by simple stimulus-response relations, and where changes to the model variables (e.g., body temperature and Mu intero) can be caused by other internal variables (e.g., the interoceptive prior). Unless the effects of these other variables are measured or inferred using a computational model, the functioning of the system will be difficult to understand^2^; see also [48] for a broader discussion of the limitations of treating the brain as a purely input-output system.

### Third simulation: failures of interoceptive control and the importance of precision

The first two (homeostatic and allostatic) simulations of interoceptive inference have illustrated how an Active Inference framework can be used to model problems of biological regulation. Notably, in these simulations, both homeostatic and allostatic interoceptive control required precise (i.e., low variance) interoceptive streams. Here, we remind that precision is a technical term that refers to the inverse variance of the probability distributions included in the generative model. The precision of sensory (exteroceptive and interoceptive) channels is normally assumed to be proportional to signal-to-noise ratio, meaning that more (less) informative signals should be weighted more (less) during the inference. However, it has been proposed that some psychopathological conditions are characterized by interoceptive insensitivity, or the failure to sense salient interoceptive changes, which may be caused by aberrant precision weighting during interoceptive inference [49].

In the third simulation, we illustrate the maladaptive effects of setting the precision of interoceptive streams to excessively low levels. In full formulations of Active Inference, precision is a parameter that can be inferred (in this case the corresponding prior beliefs are known as ‘precision expectations’). However, in this simulation, we do not allow the model to infer values of interoceptive precision; rather, we explicitly set the interoceptive precision to a very low value (here, 0.01) to render the model largely insensitive to interoceptive changes.

Results from this simulation are shown in Figure 4. The setup and generative model are identical to the second simulation. However, in this simulation we decrease the precision of predictions about interoceptive sensory data, which in turn has two effects. First, it causes posterior beliefs about interoceptive data to be biased towards prior beliefs, as the reduced precision downweights the influence of sensory data. Second, it reduces the influence of autonomic reflexes, which are driven by the magnitude of precision-weighted prediction error.

**Figure 4.**
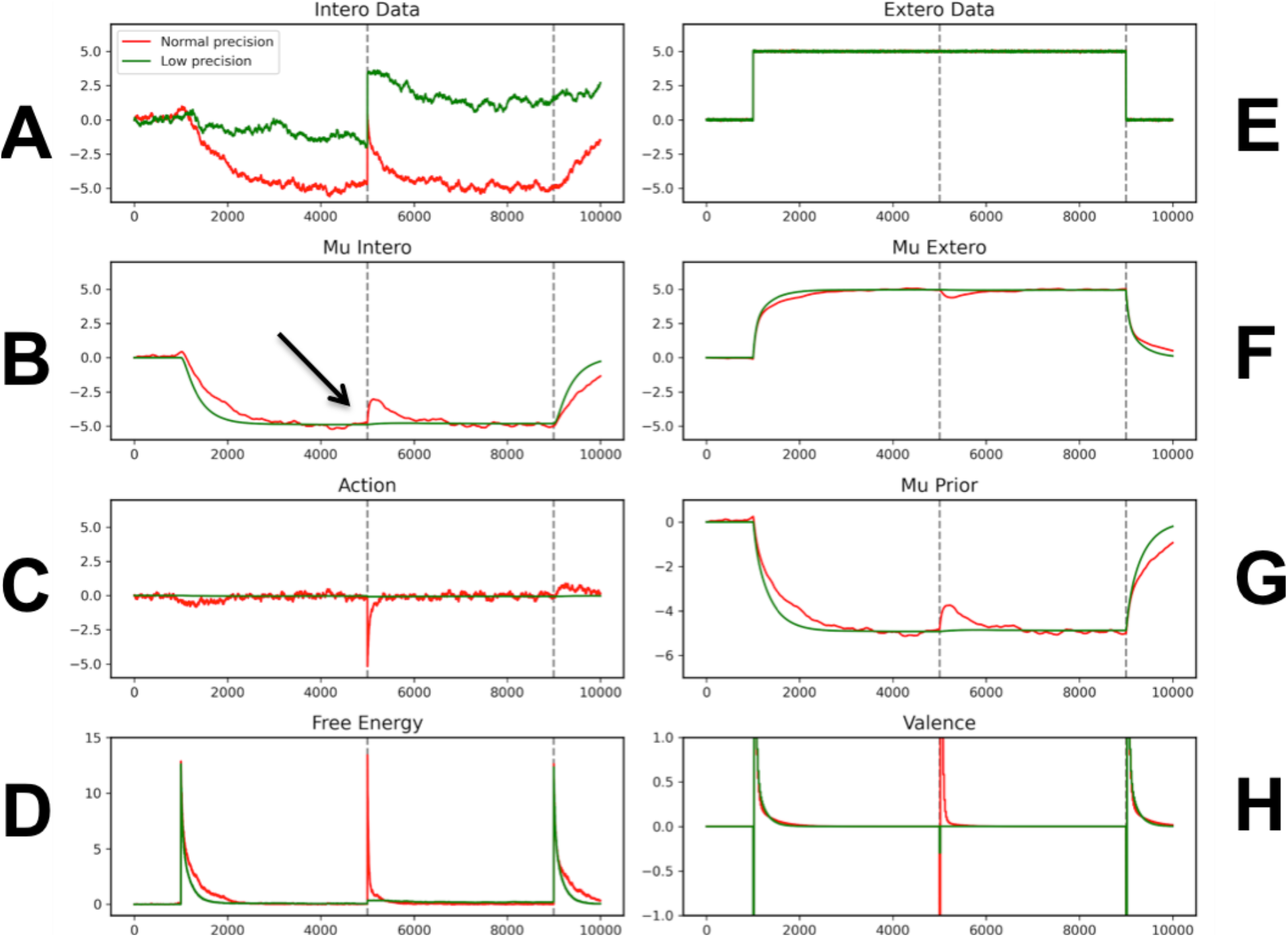
Results of the third simulation, comparing the case the precision of interoceptive streams is set to a low level (0.01, green line) versus the case where precision is within normal range (1, red lines). The latter case is identical to Figure 3.

The reduced influence of sensory data on the posterior belief can be appreciated by noticing that despite the interoceptive changes (Figure 4A), the Mu intero variable does not significantly change (see the arrow in Figure 4B, green line) and remains almost identical to the prior (Figure 4G). The reduced influence of autonomic reflexes can be appreciated by considering that compared to the first two simulations, the magnitude of autonomic actions (Figure 4C, green line) is very small. In turn, this causes a maladaptive interoceptive control, which becomes apparent by comparing much larger increase in actual body temperature under low precision (Figure 4A, green line) than that observed with normal precision (Figure 4A, red line).

This third simulation illustrates that in the absence of precise interoceptive streams, interoceptive changes are not sensed, leading to a form of interoceptive insensitivity. Furthermore, and most critically, in virtue of suboptimal precision weighting, the model fails to generate appropriate corrective responses. There is an increasing consensus that the aberrant precision weighting of exteroceptive, proprioceptive and/or interoceptive streams may be an important factor in the explanation of various psychopathological conditions, such as psychosis, chronic fatigue, depression, eating disorders and panic disorders [23,24,26–30,49]. While some theoretical and computational models have explored these implications by examining the effects of low precision on beliefs and perception (using predictive coding), only few of them explicitly incorporated control (using Active Inference). Our simulation adds to this prior work by illustrating how imprecise interoceptive streams can have maladaptive effects on both perception and control. The maladaptive effects of low precision interoceptive streams on perception are evident in the Mu intero variable, which are not updated in the light of novel (interoceptive) data. These effects would also be observed in a more restricted (predictive coding) model. The maladaptive effects of low precise interoceptive streams on control are evident in the fact that autonomic responses are not sufficient to cancel out interoceptive problems, causing body temperature to arise significantly (compared to the second simulation). These effects of low precise interoceptive streams on control are more subtle and can only be simulated by using a full Active Inference (or similar) model that includes action dynamics.

### Fourth simulation: goal-directed interoceptive control

The fourth simulation illustrates a more sophisticated case of predictive control: the case of a runner who has to decide whether to bring a bottle of water before a long run. As in the second simulation, the runner predicts a future increase of her body temperature; but this predicted increase is now so large that it cannot be compensated by autonomic reflexes alone (e.g., vasodilatation by itself cannot compensate for the large increase of body temperature caused by a long run). Hence, the runner has to form a goal-directed plan to bring a bottle of water, in anticipation of his or her future needs. We call this example of interoceptive control “goal-directed” (rather than allostatic) to highlight the fact that (unlike the previous simulations) it requires considering actions courses or policies (in a POMDP setting) and to form explicit predictions about the future values of exteroceptive and interoceptive states.

The fourth simulation models this scenario by extending the generative model used in the second simulation, to render it both hierarchical (with the inclusion of two levels) and temporally deep (with the inclusion of variables that represent future states), see Figures 5 and 6. Similar to the model used in the second simulation, the model used for this simulation includes a continuous component at the lower hierarchical level, which interfaces with interoceptive sensory data. However, the set point (prior probability) for this model is not fixed, but is instead set dynamically by a hierarchically superordinate model. This hierarchically superordinate model is composed of discrete variables which assimilate multi-modal observations, from both the continuous interoceptive model (whose highest-level acts as an observation for the lowest level of the discrete model), and from exteroceptive observations which convey whether the agent is carrying water and how long they have been running. The variables in the discrete model represent the probability that the agent is a) carrying water and b) the length of time the agent has been running (here, time is discretized in 20 intervals). Importantly, these variables span representations of the present time (s_1_) and of future times (s_2_,…, s_n_), which renders the generative model temporally deep.

**Figure 5.**
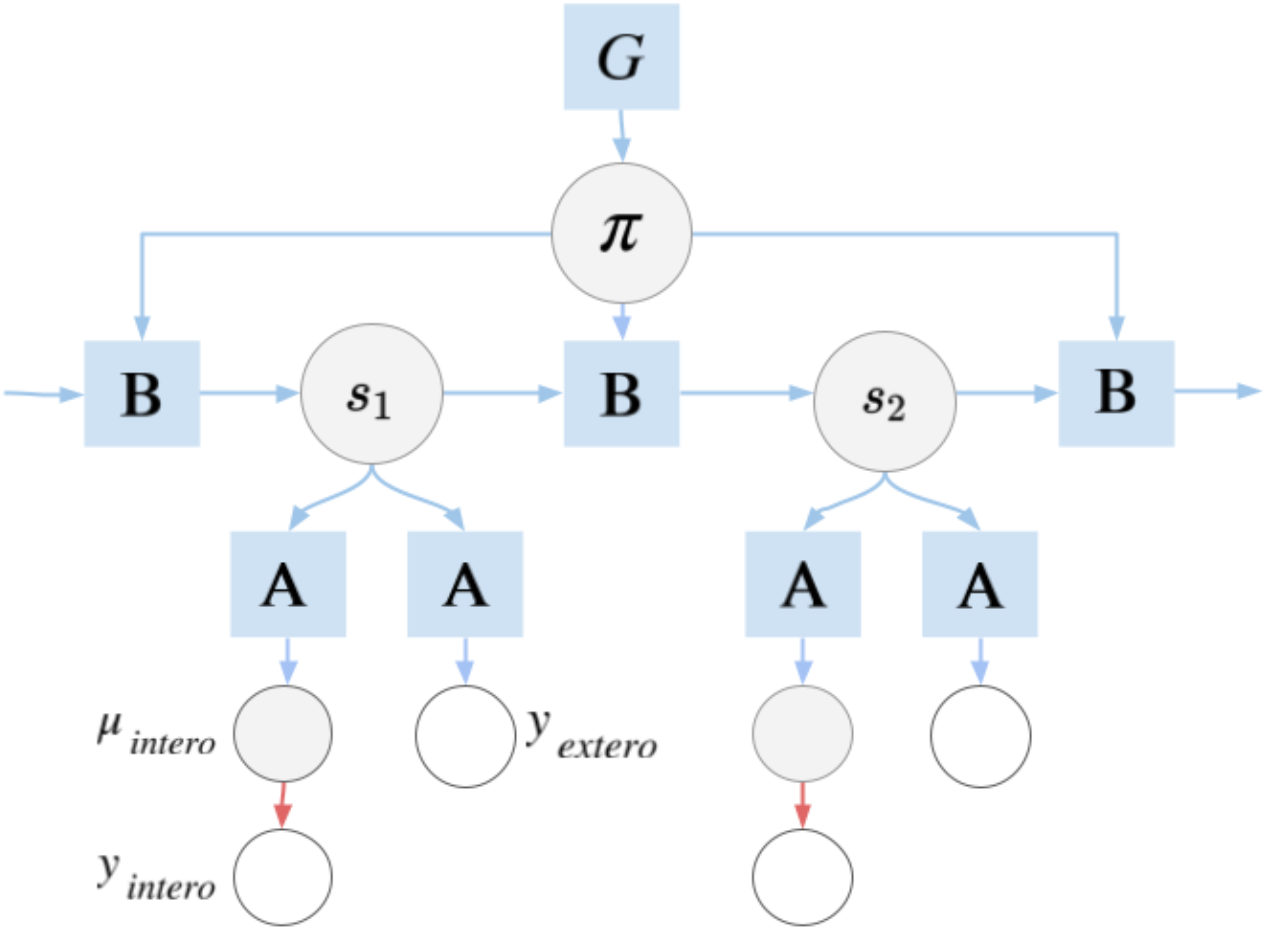
Generative model used in the fourth simulation. This generative model is hierarchical. The lower-level variables model sensory data, as in the previous simulations. The higher-level variables represent the probability that the agent is carrying water and the length of time the agent has been running (here, for simplicity, all variables are collectively called s and s1 and s2 indicate the same s at two different times). Discrete exteroceptive observations, conveying the length of time running and whether the agent is running with water, are collectively denoted *y_extero_*. Furthermore, the higher-level variable *π* denotes the two policies to run with or without water, which are illustrated schematically in Figure 6. Note that the variables of the lower level have continuous values, whereas the variables of the higher level have discrete values, which implies that the generative model is not just hierarchical but also hybrid (Karl J. Friston, Parr, et al., 2017). Finally, the model is temporally deep as the (discrete) variables of the higher level explicitly represent both the present (s_1_) and future (s_2_,…, s_n_) times. In the figure, circles indicate variables of the model. **G** indicates expected free energy; **A** indicates the likelihood function, which maps variables across hierarchical levels; **B** indicates the transition function, which maps variables across consecutive time points, conditioned on policies *π*. See the main text for explanation and (Karl J. Friston, Parr, et al., 2017) for technical details on hybrid generative models of Active Inference.

**Figure 6.**
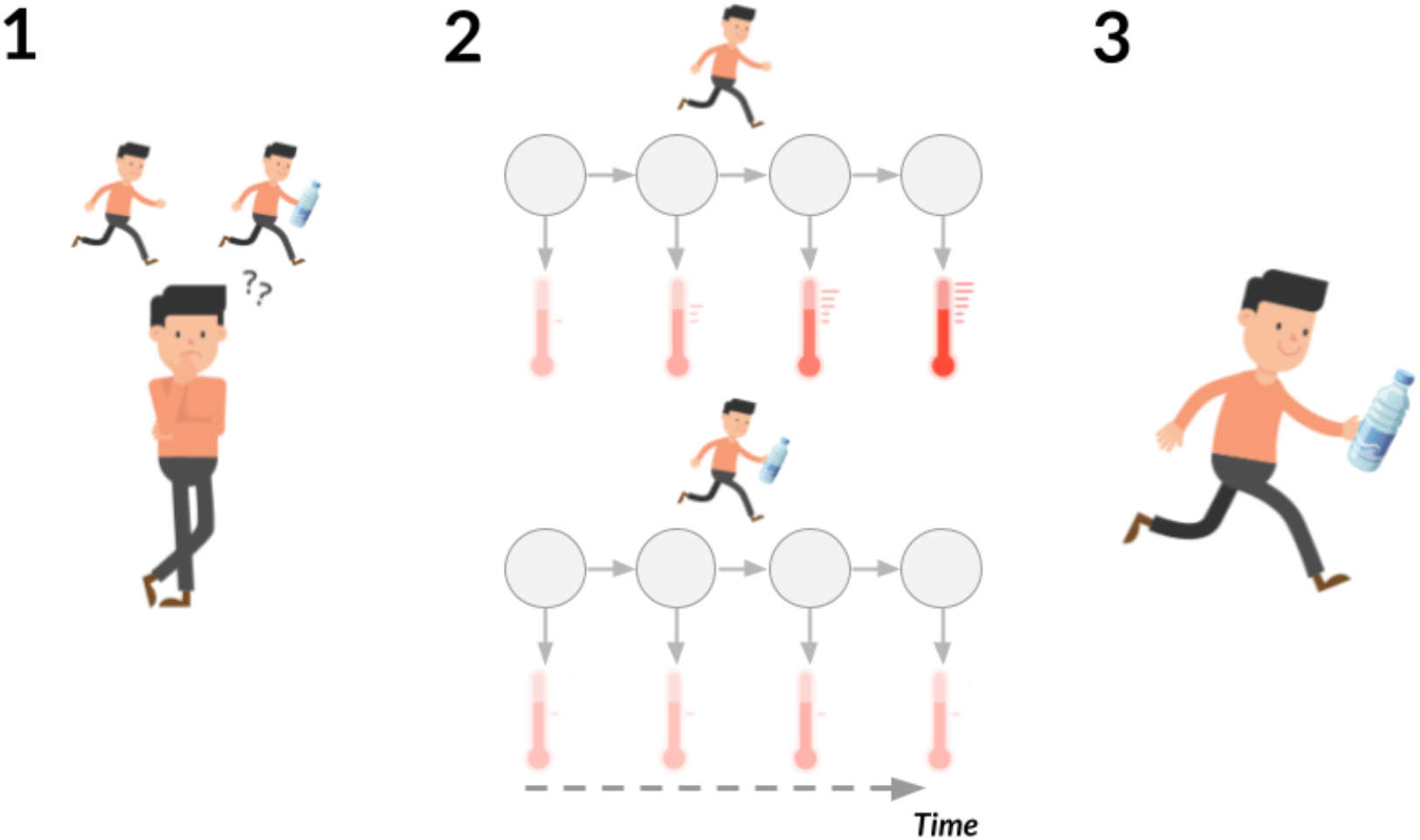
Cartoon of the (discrete) decision process of the runner, during the selection between two plans that involve bringing or not bringing a bottle of water. In step 1, the agent is faced with the decision of whether or not to bring a bottle of water while running. In step 2, the agent evaluates the future consequences of running with and without a bottle of water. Specifically, the agent predicts the consequences on body temperature as the run progresses, i.e., an increase of body temperature if the selected plan is not to bring the bottle of water (top panel) and a stable body temperature if the selected plan is to bring the bottle of water (bottom panel). In step 3, the agent chooses the preferable plan (as scored by expected free energy), which, given the prior belief that body temperature will remain constant, is running with a bottle of water.

The agent must select between one of two policies: run with water or run without water. Forming a plan to bring (or not to bring) a bottle of water requires engaging the entire generative model of Figure 5, hence encompassing both discrete and continuous variables. The planning process initially operates on higher-level (discrete) variables to predict the long-term (multimodal) consequences of bringing or not bringing water, using the transition function (called **B** in Figure 5). Then, the multimodal predictions are mapped into specific sensory (interoceptive and exteroceptive) outcomes; hence engaging the lower-level (continuous) variables, via the likelihood function (called **A** in Figure 5). The discrete model additionally encodes a prior belief that body temperature will remain at zero degrees, such that counterfactual predictions which deviate from zero will incur a prediction error (here, scored by a KL-divergence between the expected and prior beliefs about body temperature).

The logic of the policy selection process is illustrated schematically in Figure 6. As the figure shows, the agent essentially predicts two possible sequences of future states (circles) and interoceptive sensations (“thermometers”) that would follow from the selection of the two policies to run without water (top panel) and run with water (top panel). Crucially, it predicts an undesired increase of temperature under the first policy and a stable temperature under the second policy, and hence selects the latter.

The results of this fourth simulation are shown in more detail in Figure 7, which plots the multi-modal observations predicted under each policy as a function of future time. For the ‘running with water’ policy (Figure 7A), the agent predicts that it will observe itself running with water (Figure 7B), and that the amount of time spent running will increase linearly into the future (top second left panel). The agent additionally predicts its interoceptive body temperature (top second right panel), which remain roughly zero over time. Note that the predictions about running with water and the amount of time spent running use only the discrete component of the model. However, the predictions about interoceptive body temperature span both the discrete and continuous components of the model. These interoceptive predictions are facilitated by a generative model which encodes the belief that running with water will not lead to an increased body temperature.

**Figure 7.**
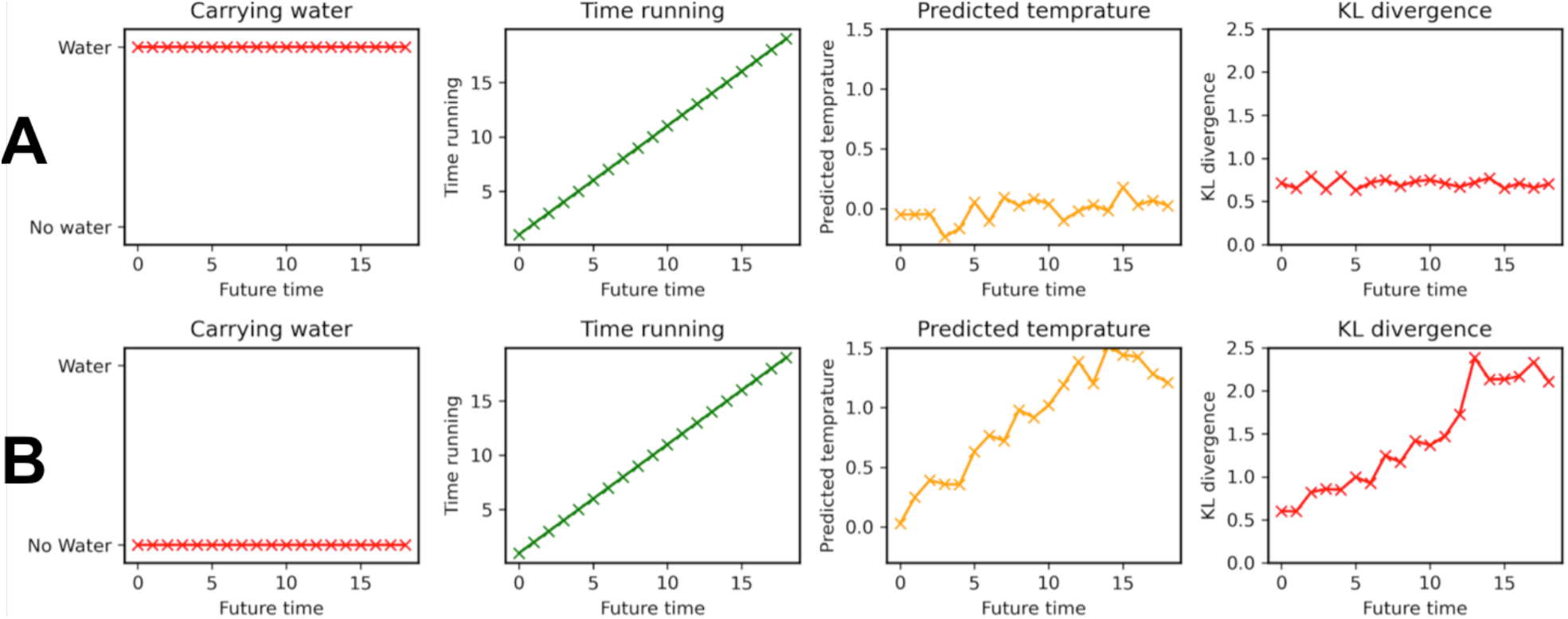
Results of the fourth simulation. In this simulation, the agent must choose between two policies – running with water (top rows) and running without water (bottom rows). Before selecting a policy, the agent predicts the future consequences of each policy, in terms of a) whether the agent will be carrying water (left column), b) how long the agent has been running (second-left column), and c) the predicted body temperature (second-right column). The predictions about body temperature incur a prediction error (or KL-divergence, right column) between what is predicted and what is expected given prior beliefs (e.g., body temperature will remain at zero). Here, higher KL-divergence indexes an inability to fulfil one’s goals (to keep body temperature around zero). Note that for simplicity, we only added uncertainty to the continuous part of the model; this is why predictions about carrying water and time running (first two panels) are perfect, but predictions about temperature (third panels) are not.

As the agent also maintains a prior belief (e.g., preference) that body temperature will remain around zero degrees, we can quantify the divergence from this prior preference using the KL-divergence between predicted and prior beliefs about body temperature (top right panel), which remains around constant for the duration of the simulation^3^. For the ‘running without water policy’, then agent predicts that it will observe itself running without water (bottom-left panel) and that the amount of time spent running will increase over time (bottom second-left panel). However, the agent now predicts that the observed body temperature will increase over time, rapidly exceeding zero and then saturating after 12 time steps. This results in the expected KL-divergence between predicted and prior beliefs increasing over time (bottom-right panel).

Given these two sets of beliefs corresponding to the policy alternatives, the agent then selects the policy with the lowest overall KL-divergence between predicted and prior beliefs about body temperature (corresponding to the expected free energy for a model without uncertainty), which is to run with water.

The key aspect of this simulation is that agent selects a policy based on *anticipated* prediction error. At the start of the simulation (time step s_1_), the agent’s body temperature is 0 (and thus accords with prior beliefs), and there is no immediate need for water. However, by engaging temporally deep hierarchical models (Figure 5), the agent is able to anticipate the future effects of action, and determine that running without water will lead to interoceptive prediction errors in the future. Here, we call this form of anticipatory regulation “goal-directed”, to distinguish it from other forms of allostatic control, such as the one shown in the second simulation, which do not require temporally deep models and explicit representations of the future. Clearly, this form of anticipatory regulation is (at least in principle) more flexible and powerful, but it also requires a more sophisticated generative model and more cognitive resources to be deployed [50].

## Discussion

The adaptive regulation of bodily and interoceptive variables, such as body temperature, water balance and glucose levels, is a fundamental challenge for any biological organism. These control problems involve (at the peripheral level) a wide array of interoceptive signals from the interior of the body and (at the central level) a sophisticated network of brain areas, broadly termed the allostatic / interoceptive network [2,3,51–53]. Besides the regulation of crucial bodily variables, the allostatic / interoceptive network is deeply involved in fundamental aspects of our emotional, motivational, self-awareness and cognitive processes [25,54–56]. Furthermore, it is becoming increasingly apparent that a malfunctioning of this network may underlie a range of psychopathological conditions, including depression, somatic symptoms, eating disorders, and possibly many other [26,29,31,57,58].

In this article, we started from the premise that living organisms address these challenges pf physiological regulation using different mechanisms. Such mechanisms range from simply reacting to sensed interoceptive errors or by anticipating them; and by only engaging autonomic actions (e.g., vasodilatation) or also performing external, goal-directed actions (e.g., drinking or finding a shaded place). This diversity is extremely useful for biological organisms, which can select the most appropriate type and level of adaptive regulation, by balancing costs (e.g., time and metabolic demands) and benefits (e.g., more or less flexibility), in a context dependent manner. However, these different forms of adaptive regulation can be difficult to disentangle - both conceptually and empirically.

Motivated by the need to clearly articulate the mechanisms and consequences of different forms of physiological regulation, we adopted a formal approach to describing mechanisms of adaptive regulation in terms of generative models of Active Inference. We used this framework to define four models of different levels of complexity. These models were illustrated in four simulations, in which the underlying generative models had different characteristics (e.g., uni- or multi-modal, shallow or deep) to address increasingly more complex (homeostatic, allostatic and goal-directed) problems of interoceptive control - and some of its possible dysfunctions.

The generative models discussed in this paper correspond to specific hypotheses about the functioning of the allostatic / interoceptive system, at both peripheral and central levels. For example, the generative model shown in Figure 1A corresponds to the hypothesis that the brain has an interoceptive schema [59]: a central representation (Mu intero) of interoceptive variables like body temperature, along with prior beliefs or “set points” for these variables (Mu prior). The generative model shown in Figure 2A incorporates the additional hypothesis that the prior (interoceptive) beliefs can be modified by exteroceptive streams. Finally, the generative model shown in Figure 5 incorporates the additional hypothesis that living organisms can form counterfactual predictions about the ways their actions will affect their future interoceptive streams.

These generative models afford different (simpler to more complex) forms of interoceptive control. Simulation 1 modeled simple reactive control in which interoceptive prediction errors are resolved by autonomic reflexes. Simulation 2 incorporated allostatic (anticipatory) control by coupling beliefs about exteroceptive signals to beliefs about future interoceptive states. Simulation 3 explored the effects of suboptimally low precision weighting on interoceptive signals, showing that effective regulation was compromised. Finally, simulation 4 implemented a hybrid scheme which integrated discrete beliefs about action policies with the continuous predictive coding schemes in the previous simulations, enabling goal-directed policy selection based on temporally deep (and counterfactual) predictions.

These simulations are not intended to model the specific details of any particular empirical situation, such as thermoregulation. Rather, we have developed these models in order to showcase the ability of the Active Inference framework to capture essential features of physiological regulation at different levels of complexity, and to make predictions about what sorts of physiological signals (and phenomenological counterparts, such as emotional valence) may be observed.

The models and simulations we have described provide a basis for formulating computationally-guided, quantitative predictions about the physiological and brain signals that one may expect to observe during experimental studies of interoception (e.g., when we raise a person’s body temperature in predictable or unpredictable ways), as well as the dysfunctions of interoceptive processing (e.g., by setting model parameters, such as precision, to incorrect values). However, to leverage the models for empirical research, it is first necessary to establish a correspondence between the time series shown in our simulations and the physiological and brain signals that we can measure during experimental studies. To establish this correspondence, it is useful to treat the variables of the generative models not just as abstract mathematical entities but (at least in principle) as relevant bodily or brain states [8]. Roughly speaking, there is a fundamental difference between the variables denoted as inputs (y) and as hidden variables (Mu) in the generative model. Interoceptive inputs and autonomic responses should putatively correspond to peripheral signals (e.g., skin temperature and conductance), whereas “Mu intero”, “Mu prior” and “free energy” signals are more likely to correspond to brain variables within the allostatic / interoceptive network, which comprises at minimum the dorsal anterior cingulate cortex (dACC), pregenual anterior cingulate cortex (pACC), subgenual anterior cingulate cortex (sgACC), and the ventral anterior insula (vaIns), see [52].

Having hypothesised these connections between variables in the model and in (the bodies and the brains of) living organisms, our models can be used to generate quantitative predictions about possible interoceptive manipulations. As Figures 2 and 3 illustrate, there are lawful relations between the intensity and duration of interoceptive inputs, internal (free energy) error responses and autonomic activity that is elicited to cancel out the errors. However, these relations are not trivial to predict without an explicit computational model. As we discussed in the Results section, this is especially true when changes of the prior and autonomic actions are triggered internally (as in the case of allostatic control) and not by external stimuli (as in the case of homeostatic control). Furthermore, without an explicit computational model, it is more difficult to distinguish causal dependencies between variables from (mere) correlations among signals. Developing explicit computational models, such as the ones discussed here, may be a valid support to the experimental analysis of allostatic / interoceptive system and the intricate dependencies between peripheral and central signals [17,27–32].

Besides the potential to support empirical investigations, computational modelling can help advancing our theoretical understanding of interoceptive inference and its dysfunctions. For example, comparing the first and second simulations sheds light on the differences between reactive (homeostatic) and predictive (allostatic) forms of control. In reactive control, priors or set points have fixed values [60], whereas in predictive control priors can be regulated proactively [5]. This becomes evident if one considers that the value of the prior in the first simulation is fixed, whereas in the second simulation it changes as a consequence of an exteroceptive event. By proactively changing priors, allostatic control permits dealing more efficiently with (predicted) interoceptive prediction errors, hence avoiding dysregulations and an “allostatic load” that can cause psychopathological conditions [26]. This becomes evident by noticing that the increase of free energy is higher in Figure 2 compared to Figure 3. This example illustrates that preparing to deal with (predicted) interoceptive prediction errors can be more efficacious than simply reacted to sensed interoceptive prediction errors.

Furthermore, our simulation of interoceptive dysfunctions (simulation 3) shows that failing to set an adaptive level of interoceptive precision can severely disrupt interoceptive control. Active Inference normally permits optimizing precision values as part of the normal inferential dynamics; but there may be pathological cases in which the precision-weighting mechanism is disrupted. To exemplify this, we simulated the effects of setting the precision of interoceptive streams to extremely low values. In this case, interoceptive changes are not sufficiently sensed and the model fails to generate appropriate corrective responses. This is a form of interoceptive insensitivity that can be observed in various psychopathologies [49]. In keeping, previous theoretical and computational studies have highlighted that aberrant precision settings can cause various perceptual problems, such as hallucinations [23,61] and the reporting of false symptoms [58], which are often associated to psychopathological conditions. However, how our third simulation exemplifies, the psychopathological consequences of incorrect precision settings can go beyond perception to cause dysfunctions of interoceptive and allostatic regulation - which in turn can cause allostatic load (as observed in depression) [26] or maladaptive compensatory strategies (as observed in anorexia) [24]. There is increasing consensus that interoceptive and allostatic disorders may be crucial for the genesis of several psychopathological conditions [2,11,30,56,57]. Designing and validating novel models of (adaptive and maladaptive) interoceptive regulation can help shed light on psychopathological conditions, by advancing our conceptual understanding and by providing sophisticated tools for quantitative hypothesis testing.

## Funding

GP was funded by the European Union’s Horizon 2020 Framework Programme for Research and Innovation under the Specific Grant Agreement No. 945539 (Human Brain Project SGA3) and the European Research Council under the Grant Agreement No. 820213 (ThinkAhead). AT and AKS are grateful to the Dr. Mortimer and Theresa Sackler Foundation, which supports the Sackler Centre for Consciousness Science. CLB is supported by BBRSC grant number BB/P022197/1.

## Footnotes

1 The mapping between exteroceptive variables and interoceptive priors does not need to be always inverse. In case the generative model includes a prior over the amount of metabolic resources (e.g., oxygen) to be consumed, this prior should plausibly increase before a run, to engage autonomic actions (e.g., hyperventilation) that increase metabolic resources. Furthermore, we could have designed an alternative generative model for thermoregulation, in which the Mu exteroceptive and Mu prior variables were not (inversely) related. In this alternative generative model, Mu exteroceptive would be coupled with a novel variable (a belief that temperature will rise) and in turn this novel variable would be (inversely) related to the Mu prior. These examples show that there are various ways to design generative models for interoceptive and autonomic control, which can be validated empirically.

2 Note that we described our simulation as if there were a specific exteroceptive (e.g., visual) stimulus that triggered internal adjustments of the prior and autonomic responses. However, in practical condition, this is hardly the case. Following our example, there may not be a specific sensory stimulus that signals the beginning of the mobilization of metabolic resources in athletes.

3 Note that for simplicity, here we used KL-divergence rather than expected free energy, as the latter is equivalent to the former if (as in our simulations) there is no ambiguity about the current state.

## Notes

### Competing Interest Statement

The authors have declared no competing interest.

